# Characterization and engineering of a blue-sensitive, Gi/o-biased, and bistable ciliary opsin from a fan worm

**DOI:** 10.1101/2024.11.21.624670

**Authors:** Sachiko Fukuzawa, Tomoki Kawaguchi, Takushi Shimomura, Yoshihiro Kubo, Hisao Tsukamoto

**Author notes:** Correspondence should be addressed to Hisao Tsukamoto; Department of Biology, Graduate School of Science, Kobe University, 1-1, Rokkodai-cho, Nada-Ku, Kobe, 657-8501, Japan.

## Abstract

Ciliary opsins have been identified not only in vertebrates but also in invertebrates. An invertebrate ciliary opsin was recently identified in the fan worm *Acromegalomma interruptum* (formerly named *Megalomma interrupta*); however, its spectral and signaling characteristics are unknown. In the present study, we characterized the spectral properties and light-induced cellular signaling properties of the opsin (*Acr*InvC-opsin). *Acr*InvC-opsin showed an absorption maximum at 464 nm and upon blue-light absorption, the spectrum was red-shifted by approximately 50 nm. The two states are inter-convertible by illumination with blue and orange light. Blue light illumination of *Acr*InvC-opsin caused specific coupling with Gi, sustained Gi dissociation, decreased intracellular cAMP levels, and activation of GIRK channels. The cellular responses by the activated opsin were partially terminated by orange light illumination. These light-dependent responses indicate that the InvC-opsin is a typical bistable pigment wherein the resting and activated states can be inter-converted by visible light illumination. We also attempted to modulate the spectral and functional properties of *Acr*InvC-opsin using site-directed mutagenesis. Substitution of Ser-94 with Ala caused little spectral shift in the resting state but a further red-shift of ∼10 nm in the activated state, indicating that the absorption spectra of the two states were tuned differently. In contrast, the S94A substitution did not significantly affect the light-dependent signaling properties of *Acr*InvC-opsin. Because *Acr*InvC-opsin is a blue-sensitive, Gi/o-biased, and bistable pigment, it has the potential to serve as an optical control tool to specifically and reversibly regulate Gi/o-dependent signaling pathways by visible light.

## Introduction

Animal opsins receive and transmit external light signals in various photoreceptor cells. Photoreception in opsins is triggered by *cis* to *trans* photoisomerization of the retinal chromophore, leading to conversion of the proteins from the resting state to the activated state^1^. Thousands of opsin genes have been identified in a wide variety of animals, and are classified into several subgroups based on their amino acid sequence similarity^2,3^. One simple classification divides the opsins into two subgroups: ciliary and rhabdomeric opsin groups^4^. The ciliary opsin (c-opsin) group mainly contains Gi/o-coupled opsins in the ciliary photoreceptor cells, whereas the rhabdomeric opsin (r-opsin) group mainly contains Gq-coupled opsins in the rhabdomeric photoreceptor cells. Vertebrates (deuterostomes) and invertebrates (protostomes) had been thought to exclusively possess ciliary and rhabdomeric opsins, respectively, but many vertebrates and invertebrates have been proved to possess both ciliary and rhabdomeric opsins^3^.

The vertebrate visual pigment is the most well-known and well-studied ciliary opsin whose activated state is thermally unstable and spontaneously releases the isomerized (all-*trans*-) retinal^5^. In contrast, some invertebrate ciliary opsins (InvC-opsins), as well as some vertebrate ciliary opsins and many rhabdomeric opsins, form stable activated states that hold all-*trans*-retinal, and they can be re-converted to the resting (dark) state via *trans*-*cis* (re)isomerization of the retinal^2,3^. The opsins having such biochemical and photochemical characteristics are called as the “bistable” pigments^6^. To completely deactivate a bistable opsin by light, sufficient separation between the absorption spectra of the resting (deactivated) and activated states is required to enable selective illumination of the activated state. For example, a bistable ciliary opsin in mosquito (mosquito Opn3) shows highly overlapped absorption spectra of the resting and activated states, and illumination with any wavelength of light produces a mixture of the resting and activated states^7^. In contrast, lamprey parapinopsin, a vertebrate bistable ciliary opsin, shows a large (∼130 nm) spectral shift upon activation from the UV region to the green region, and green light illumination can effectively and completely deactivate this opsin^6,8^.

Extensive studies on InvC-opsins in annelids, particularly the marine ragworm *Platynereis dumerilii,* have revealed their spectral properties and physiological functions. *Platynereis* c-opsin1 and c-opsin2 are UV- and blue-sensitive bistable pigments, respectively, both of which play different roles in photoreceptive functions of the animal^9–12^. Our recent study on the c-opsin1 identified non-canonical features as a bistable pigment^13^. Similar to typical bistable opsin, the c-opsin1 is inter-convertible between the resting and activated states by UV and yellow light illumination; however, the activated state is spontaneously inactivated in a time-dependent (light-independent) manner^13^. *Platynereis* c-opsin1 and lamprey parapinopsin have been established as useful optogenetic tools to turn ON and OFF the Gi/o-dependent cellular responses by different colors of light^8,14,15^. *Platynereis* c-opsin2 is a typical bistable pigment that forms a stable activated state, and the absorption spectra of the resting and activated states are somewhat overlapped^9^. These studies indicate that spectral and signaling characteristics of InvC-opsins are diversified require experimental assessment.

A recent study revealed that another annelid, the fan worm *Acromegalomma interruptum* (originally named *Megalomma interrupta*), expresses an InvC-opsin (hereafter referred to as *Acr*InvC-opsin; Supplemental Fig. S1) in the radiolar eyes^16^. This study suggests that *Acr*InvC-opsin drives the Gi/o-dependent signaling cascades in a light-dependent manner based on transcriptome and molecular phylogenetic profiles^16^; however, the spectral and signaling properties of *Acr*InvC-opsin have not been experimentally investigated.

In the present study, we characterized the spectral and signaling characteristics of *Acr*InvC-opsin. This opsin is blue-sensitive, and upon blue light absorption, it is activated and red-shifted by approximately 50 nm. The activated state was re-converted to the resting state upon orange light absorption. Activated *Acr*InvC-opsin is predominantly coupled with Gi, and causes sustained Gi dissociation, leading to Gi/o-dependent cAMP reduction and GIRK activation. Moreover, substitution of the serine residue at position 94 with alanine caused a ∼10 nm red-shift in the activated state. Thus, *Acr*InvC-opsin is a blue-sensitive, Gi/o-coupled, and bistable pigment, and its molecular characteristics can be modulated by site-directed mutagenesis. We discuss the functional diversity of invertebrate ciliary opsins and possibility of engineering *Acr*InvC-opsin as a Gi/o-biased and bistable optogenetic tool.

## Materials and Methods

### Ethics statement

All animal experiments in this study were approved by the Animal Care Committee of the National Institutes of Natural Sciences (an umbrella institution of National Institute for Physiological Sciences, Japan), and were performed in accordance with its guidelines.

### Construction of WT and S94A mutant for heterologous expression in mammalian cultured cells and Xenopus oocytes

The cDNA of *Acr*InvC-opsin (MF145115.1)^16^ was optimized for human codon usage and synthesized by Eurofins Genomics, Japan Inc. The synthesized cDNA was fused with the coding sequence of 36 amino acids on N-terminus of *Platynereis* c-opsin1^4^ and the coding sequence of 1D4 tag (ETSQVAPA), which is a recognition sequence of the antibody 1D4, followed by the truncation of C-terminus (see Fig. 1A and Supplemental Fig. S2A). The cDNAs of *Acr*InvC-opsin WT and the S94A mutant were inserted into the *Eco*RI/*Not*I site in an expression vector pMT or *EcoRI/HindIII* site in an expression vector pGEMHE using in-Fusion HD (TAKARA, Japan). The cDNAs of Lg-BiT inserted mouse Giα_2_, human Gβ_1_, and the Sm-BiT fused human Gγ_2_ (C68S) were constructed and inserted into the pMT vector according as described previously^13^.

**Fig. 1.**
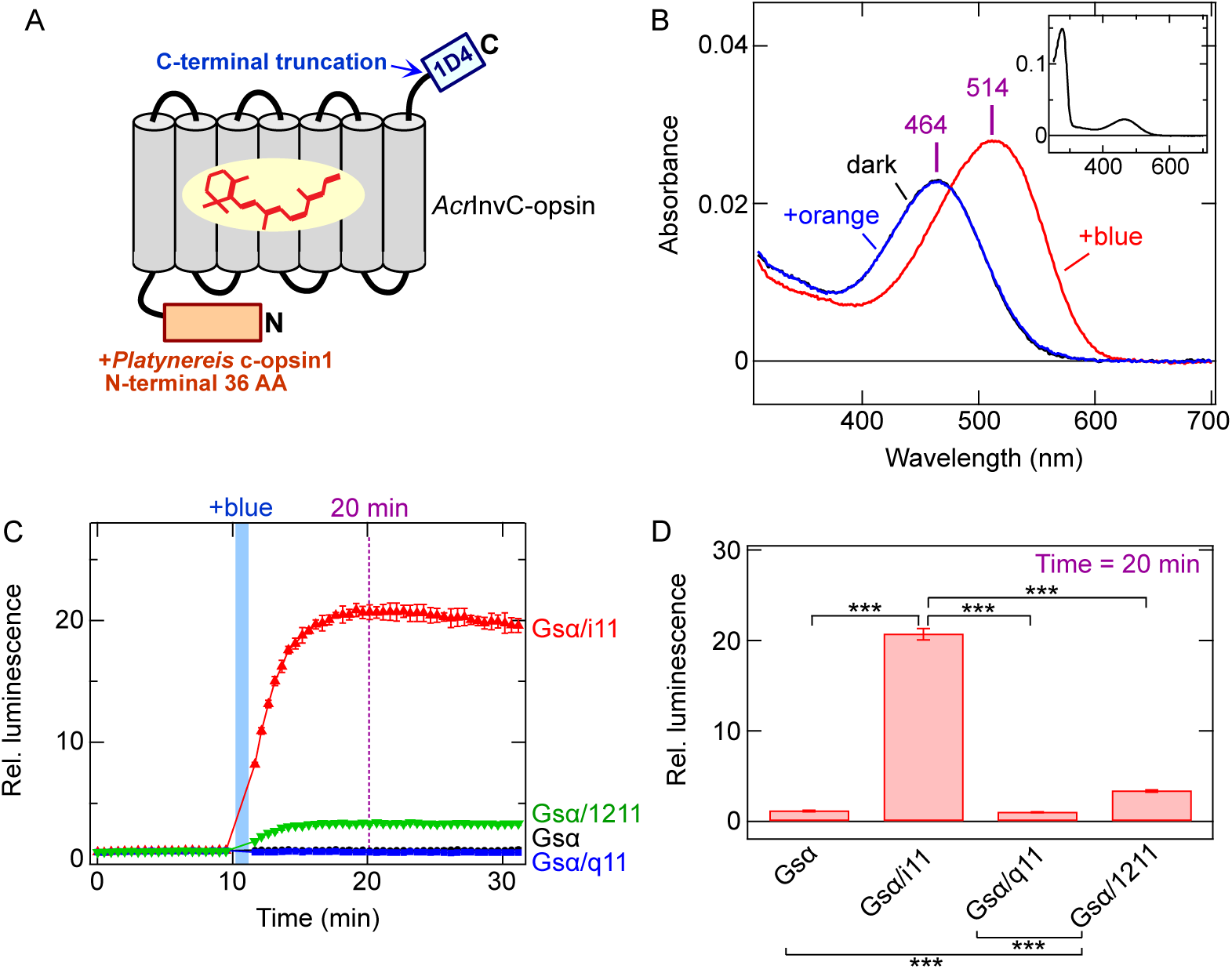
Construct, spectral properties, and G protein-coupling specificity of *Acr*InvC-opsin. *A*, Schematic drawing to describe the construct of *Acr*InvC-opsin used in this study. In the construct, 36 amino acids of N-terminus in *Platynereis* c-opsin1 was fused, and C-terminal sequence was removed, and the 1D4 sequence (ETSQVAPA) was added to the opsin. See Supplemental Fig. S2A for detail. *B*, Absorption spectra of purified *Acr*InvC-opsin. Spectra in the dark (*black*), after blue illumination (*red*), and after subsequent orange light (>580 nm) illumination (*blue*) are shown. The *black* and *blue* spectra are almost superimposed (but see Supplemental Fig. S3), indicating that orange light is primarily absorbed by the red-shift photoproduct. In the dark, α_max_ value is 464 nm, and after blue light illumination, the value is shifted to 514 nm. (*Inset*) Absorption spectrum of *Acr*InvC-opsin in the dark in a wider wavelength range. *C*, GsX GloSensor assay profiles for *Acr*InvC-opsin. Blue light induced changes in cAMP biosensor (GloSensor) luminescence in COS-1 cells expressing Gsα (*black*), Gsα/i11 (*red*), Gsα/q11 (*blue*), or Gsα/1211 (*green*) with the InvC-opsin are plotted. Luminescence levels are normalized to the value at the starting point (time = 0 min). Error bars indicate the S.D. values (n = 3). Blue bar shows blue light illumination and the duration of illumination are also indicated. *D*, Comparison of relative luminescence levels at 20 min (strictly, 20.14 min) in panel *C*. Error bars indicate the S.D. values (n = 3). The value of “Gsα/i11” is significantly different from the values of “Gsα”, “Gsα/q11”, and “Gsα/1211” (*P* < 0.001***). The values of “Gsα/1211” are significantly different from the values of “Gsα” and “Gsα/q11” (*P* < 0.001***). The statistical differences were evaluated using Tukey’s test following one-way ANOVA.

### Protein expression and purification of AcrInvC-opsin

*Acr*InvC-opsin WT and S94A mutant were transiently expressed in COS-1 cells (10 plates), and the cells were harvested 48 h after transfection as described previously^10,17^. The collected cells were incubated with 11-*cis*-retinal overnight, and membrane proteins were solubilized with 1.25 % DDM (Dojindo, Japan), 20 mM HEPES, 140 mM NaCl, 0.25 % cholesterol hemisuccinate (Sigma-Aldrich, St. Louis, MO) 25 mM Tris, 10 % glycerol, pH 7.0. The solubilized materials were mixed with 1D4-agarose overnight, and the mixture was transferred into Bio-Spin columns (Bio-rad, Hercules, CA). The columns were washed with 0.05 % DDM, 2 mM ATP, 1 M NaCl, 3 mM MgCl_2_, 0.01 % cholesterol hemisuccinate, 1 mM Tris, 10 % glycerol in PBS, and subsequently washed with 0.05 % DDM, 140 mM NaCl, 0.01 % cholesterol hemisuccinate, 1 mM Tris, 10 % glycerol, 20 mM HEPES, pH 7 (buffer A). The 1D4 tagged pigments were eluted with buffer A containing 0.45 mg/mL 1D4 peptide (TETSQVAPA) (TOYOBO, Japan).

### UV-Vis spectroscopy and photoreaction of AcrInvC-opsin

Absorption spectra of purified opsins were recorded with a Shimadzu UV-2600 spectrophotometer (Shimadzu, Japan). The samples were kept at 10 °C. An optical interference filter VPF-50S-10-45-44000 (Sigma-Koki, Japan), which transmits light around 440-nm, was used for illumination of *Acr*InvC-opsin with blue light (duration: 1 min, light intensity: ∼30 mW/cm^2^, light source: 250 W halogen lamp). In order to convert the photoproduct to the original state, a longpass filter SCF-50S-58O (Sigma-Koki) was used (duration: 30 sec, light intensity: ∼140 mW/cm^2^, light source: 250 W halogen lamp) as orange light.

### Transfection to COS-1 cells for GloSensor and NanoBiT G protein dissociation assays

Opsins, GloSensor assay sensor (coded by pGlo-22F), and G proteins (with or without NanoBiT-tag) were transiently expressed in COS-1 cells using polyethyleneimine as described previously^13^. For the GloSensor assay, each well of 96-well assay plate (Corning, Kennebunk, ME) was transfected with 50 ng *Acr*InvC-opsin plasmid (or 5 ng or 0.5 ng, see Figs. 3C and 4H), 50 ng pGlo-22F plasmid (Promega, Madison, WI), and 500 ng polyethyleneimine in 25 μL Opti-mem (Gibco, Waltham, MA) and 75 μL D-MEM (Wako, Japan) containing 10 % (v/v) FBS, 100 units/mL penicillin, and 100 μg/mL streptomycin. For the GsX Glosensor assay^14,18^, 16.7 ng plasmid coding Gsα, Gsα/i11, Gsα/q11, or Gsα/1211 was also transfected. For the NanoBiT G protein dissociation assay, each well of 96-well assay plate (Corning) was transfected with 50 ng *Acr*InvC-opsin plasmid, 8.5 ng Lg-BiT inserted Giα_2_ plasmid, 40 ng Gβ_1_ plasmid, 40 ng Sm-BiT fused Gγ_2_ plasmid, and 500 ng polyethyleneimine in 25 μL Opti-mem (Gibco) and 75 μL D-MEM (Wako) containing 10 % (vol/vol) FBS, 100 units/mL penicillin, and 100 μg/mL streptomycin.

### GloSensor assay and GsX Glosensor assay

The transfected COS-1 cells were incubated at 37 °C, 5 % CO_2_ for 2 days, and the medium was aspirated, followed by addition of HBSS (145 mM NaCl, 10 mM D-glucose, 5 mM KCl, 1 mM MgCl_2_, 1.7 mM CaCl_2_, 1.5 mM NaHCO_3_, 10 mM HEPES, pH 7.4) containing 1 μM (or 100 M or 10 nM, see Figs. 3A and 4F) 11-*cis*-retinal, 2 % (vol/vol) GloSensor cAMP reagent stock solution, and 1 μM forskolin. For the GsX GloSensor assay (Fig. 1C), forskolin was not added to HBSS. Then, the cells were incubated at room temperature for ∼2 h to stabilize luminescence levels. Luminescence was measured using GM-2000 or GM-3510 microplate reader (Promega). Luminescence level of each well was measured every 30 sec with integration time of 0.9 sec. To illuminate *Acr*InvC-opsin, luminescence measurement was interrupted, the plate was ejected, and the plate was illuminated by blue or orange light. Blue and orange light sources are 460 nm (duration: 1 min, light intensity, ∼0.3 mW/cm^2^) and 600 nm (duration: 1 min, light intensity, ∼0.4 mW/cm^2^) from Opto-spectrum generator L12194 (Hamamatsu, Japan), respectively. After illumination, luminescence measurement was resumed. The measured luminescence levels were normalized to the level at the starting point (time = 0 min).

### NanoBiT G protein dissociation assay

The transfected COS-1 cells were incubated at 37 °C, 5 % CO_2_ for 1 day, and the medium was aspirated, followed by addition of HBSS containing 1 μM 11-*cis*-retinal and 5 μM coelenterazine h (Wako). Then, the cells were incubated at room temperature for ∼2 h to stabilize luminescence levels. Luminescence was measured using GM-2000 or GM-3510 microplate reader (Promega). Luminescence level of each well was measured every 30 sec with integration time of 0.3 sec. To illuminate *Acr*InvC-opsin, luminescence measurement was interrupted, the plate was ejected, and the plate was illuminated by 460 nm (duration: 1 min, light intensity, ∼0.3 mW/cm^2^) or 600 nm (duration: 1 min, light intensity, ∼0.4 mW/cm^2^) from Opto-spectrum generator L12194 (Hamamatsu). After illumination, luminescence measurement was resumed. The measured luminescence levels were normalized to the level at the starting point (time = 0 min).

### Preparation of Xenopus oocytes and cRNA injection

*Xenopus* oocytes were isolated from frogs as described previously^19–21^. Briefly, *Xenopus* oocytes were surgically collected from frogs anesthetized in water containing 0.15% tricaine and treated with 2 mg/ml collagenase (Sigma-Aldrich) for 3–4 h to remove the follicular membrane. 5’-capped cRNA was prepared from the pGEMHE vector containing *Acr*InvC-opsin cDNA (see above) using an in vitro transcription kit (mMESSAGE mMACHINE Kit, Life Technologies). Typically, we injected the *Acr*InvC-opsin cRNA (∼300 pg/oocyte) with rat GIRK1 and mouse GIRK2 cRNAs (∼25 ng/oocyte and ∼12.5 ng/oocyte, respectively) in 50 nL of water/oocyte. The oocytes injected with cRNA were incubated in the standard frog Ringer solution (MBSH), a standard frog Ringer solution (88 mM NaCl, 1 mM KCl, 0.3 mM Ca(NO_3_)_2_, 0.41 mM CaCl_2_, 0.82 mM MgSO_4_, 2.4 mM NaHCO_3_, and 15 mM HEPES (pH 7.6) with 0.1% penicillin-streptomycin solution), at 17 °C in the dark chamber for 2 days.

### Electrophysiology

We used a conventional two-electrode voltage clamp technique^20,22^ to measure photoresponses caused by opsins. Before electrophysiological recording, oocytes injected with cRNAs were incubated in the standard frog Ringer solution (MBSH) containing 1 μM 11-*cis*-retinal (1/4000 volume of 4 mM retinal in ethanol was added to MBSH) for ∼1 h at 17 °C in the dark chamber to form photosensitive pigments. All electrophysiological recordings were performed in a dark room, using only a dim red light. Light-induced electrophysiological responses were recorded in a bath solution (96 mM KCl, 3 mM MgCl_2_, and 5 mM HEPES (pH 7.4)). The tip resistance of the glass electrodes was 0.2–0.5 MΩ when filled with the pipette solution (3 M potassium acetate and 10 mM KCl). The increase in inward K^+^ current as a result of Gi/o activation by opsins was monitored by two-electrode voltage clamp technique using an OC-725C amplifier (Warner Instruments, Hamden, CT) at room temperature with continuous hyperpolarizing pulses of 0.2 s to -100 mV every 2 s from the holding potential of 0 mV and subsequent 0.2-s pluses of 40 mV. *Acr*InvC-opsin was illuminated with blue light (470 nm LED, Sarspec, Portugal; light intensity, ∼0.3 mW/cm^2^) or orange (>580 nm) light (>580 nm, a longpass filter SCF-50S-58O, Sigma-Koki with 3-W white LED light, OptoCode, Japan; light intensity, ∼10 mW/cm^2^). Data acquisition was performed by a digital converter (Digidata 1440, Molecular Devices, Sunnyvale, CA) and pCLAMP 10 software (Molecular Devices).

## Results

### Spectral properties of AcrInvC-opsin

The previous study identified an InvC-opsin gene in the fan worm, *Acromegalomma interruptum*, but the reported opsin gene sequence lacks the N-terminal sequence^16^. We constructed the *Acr*InvC-opsin with a simple initial Met residue or 36 N-terminal amino acid residues of the closely related *Platynereis* c-opsin1^4^ at the N-terminus (Fig. 1A, and Supplemental Figs. S1 and S2A). We also added the 1D4 sequence (ETSQVAPA) to the truncated C-terminus (Fig. 1A and Supplemental Fig. S2A) because truncation of the C-terminus in several InvC-opsins, such as mosquito Opn3, increased their expression levels in mammalian cultured cells^7^. The addition of the N-terminal sequence of *Platynereis* c-opsin1 increased expression level of the opsin more than the simple addition of initial Met residue (Supplemental Fig. S2B). Hereafter, the N- and C-terminal modified *Acr*InvC-opsin is named as “WT” in this study. *Acr*InvC-opsin WT was expressed in mammalian cultured COS-1 cells, mixed with 11-*cis*-retinal, extracted with detergent DDM, and purified using the 1D4 antibody-conjugated column (see “Materials and Methods”).

Purified *Acr*InvC-opsin showed an absorption maximum (αmax) at 464 nm, indicating that it is a blue-sensitive pigment (Fig. 1B). Blue light illumination converted the spectrum to a red-shifted spectrum (αmax = 514 nm), and orange light illumination of the red-shifted species re-converted the spectrum to almost the same spectrum as the original one (Fig. 1B). Strictly, very tiny fraction of the red-shifted species existed after orange light illumination (Supplemental Fig. S3), and the remaining activated species affected signaling properties (see below). These photoreactions indicate that the opsin is a typical bistable pigment in which the two stable (original and red-shifted) states are inter-convertible by illumination with different wavelengths of light.

### G protein-coupling selectivity of AcrInvC-opsin

Although the previous study concluded that *Acr*InvC-opsin is Gi/o-coupled based on phylogenetic relationships and transcriptome data^16^, the G protein-coupling selectivity of the opsin has not been experimentally tested. To examine the selectivity of G protein-coupling, we exploited a GsX GloSensor assay. This assay measures GPCR-induced increases in intracellular cAMP levels even for Gq-, Gi/o-, or G12-coupled receptors using Gsα chimeras, and provides insights into the G protein-coupling selectivity of the receptors^14,18^. We prepared Gsα mutants, in which the C-terminal 11 amino acid sequence was replaced with those of Giα (Gsα/i11), Gqα (Gsα/q11), and G12α (Gsα/1211). These Gsα chimeric mutants or WT were expressed in COS-1 cells with *Acr*InvC-opsin and a luciferase-based cAMP biosensor (see “Materials and Methods”). This assay enabled the measurement of Gs/Gi/Gq/G12 activation by the opsin as an increase in luminescence upon cAMP production. It was found that upon blue light illumination, this opsin caused a significantly larger cAMP responses with Gsα/i11 than with Gsα WT, Gsα/q11, or Gsα/1211 (Fig. 1, C and D). In addition, the opsin showed a small but significant light-dependent cAMP increase with Gsα/1211 compared with Gsα WT and Gsα/q11 (Fig. 1, C and D). These results proved that *Acr*InvC-opsin is selectively coupled with Gi/o-type G proteins, although it may only be very weakly coupled with G12. In other words, *Acr*InvC-opsin can drive Gi/o-biased signaling pathways in a light-dependent manner. We thus tested Gi/o-dependent downstream cellular responses driven by the opsin using various cellular assays and electrophysiological analyses.

### Light-induced Gi/o-dependent cellular responses driven by AcrInvC-opsin

Next we assessed the light-induced G protein activation and deactivation by *Acr*InvC-opsin. As the GsX GloSensor assays indicated *Acr*InvC-opsin is selectively coupled with Gi/o-type G proteins (Fig. 1, C and D), we tested the light-dependent dissociation and association of the Gi trimer. We also analyzed typical Gi/o-dependent cellular responses, such as Gi/oα-dependent intracellular cAMP reduction in mammalian cultured cells and Gβγ-dependent GIRK channel activation in *Xenopus* oocytes. We analyzed how these signaling responses were modulated by blue and orange light illumination of the opsin.

We assessed the dissociation and association of the Gi trimer using a NanoBiT G protein dissociation assay. In this assay, Gi dissociation is detected as a decrease in NanoLuc luminescence from the Lg-BiT fragment inserted into Giα and the Sm-BiT fragment fused with Gβγ^23^. We sequentially illuminated with orange, blue, and orange light on the COS-1 cells expressing *Acr*InvC-opsin, Lg-BiT fused Giα, Gβ, and Sm-BiT fused Gγ. We expected that the first illumination with orange light would cause no luminescence changes, the second blue light illumination would decrease the luminescence (Gi dissociation), and that the third orange light illumination would fully recover the luminescence (Gi reassociation). Obtained data showed that the first orange illumination caused a small but significant luminescence decrease, indicating somewhat Gi activation (Fig. 2A). As expected, the second blue light illumination induced further and sustained Gi activation, and the third orange light illumination canceled the second blue light-induced Gi activation (Fig. 2A). The NanoBiT assay results indicated that the species with αmax at 464 nm is the resting state and the red-shifted species is the activated state, and that the two states are inter-convertible by blue and orange light illumination (see Fig. 1B). The results also indicated that the orange light was absorbed not only by the activated state but also by the resting state because of the broadness of their absorption spectra (Supplemental Figs. S3 and S4). This is how orange light illumination caused the unintentional formation of a tiny amount of the activated state (see Supplemental Fig. S3).

**Fig. 2.**
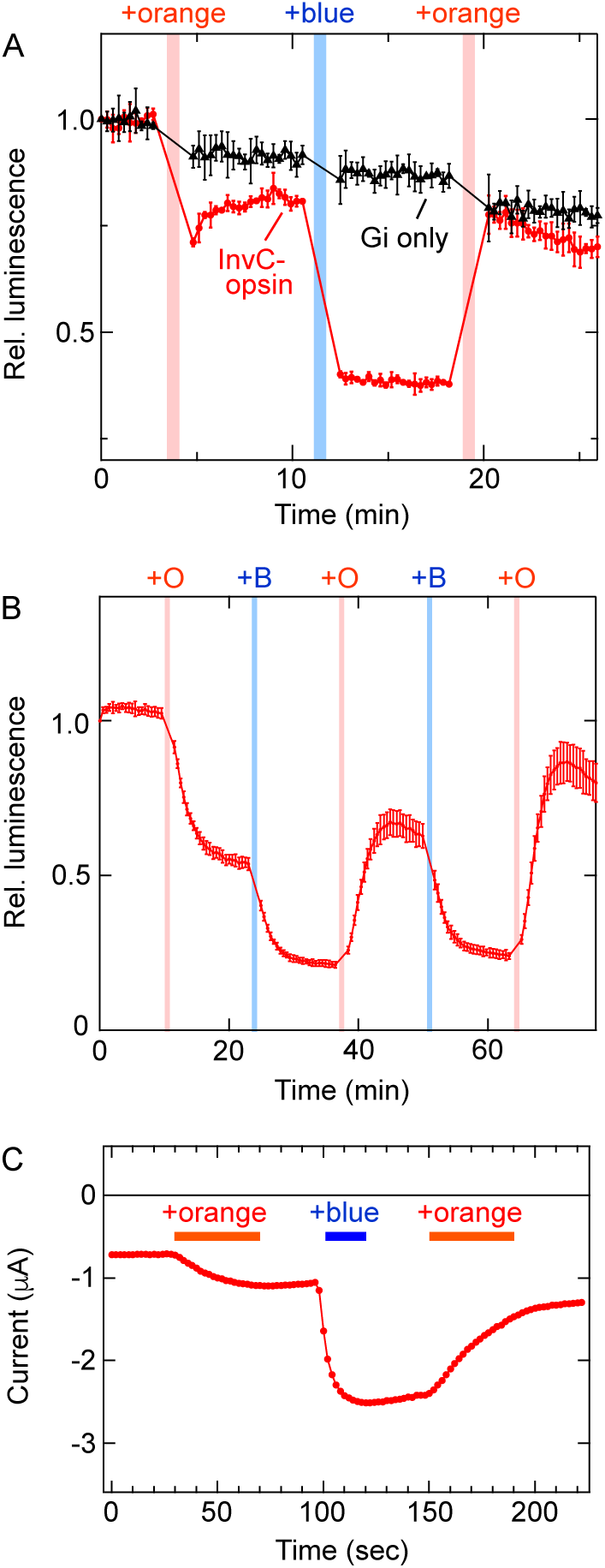
Gi dissociation and re-association, Gi/oα-dependent cAMP responses, and Gβγ-dependent GIRK responses caused by illimitation on *Acr*InvC-opsin. *A*, NanoLuc luminescence changes of NanoBiT-tagged Gi proteins upon blue and orange light illumination on *Acr*InvC-opsin WT (*red*) in COS-1 cells. The same measurement was conducted without opsins (Gi only, *black*). Error bars indicate the S.D. values (n = 3). NanoLuc luminescence levels are normalized to the value at the starting point (time = 0 min). Blue and orange bars indicate blue and orange light illuminations, respectively. *B*, Light induced changes in cAMP biosensor (GloSensor) luminescence in COS-1 cells expressing *Acr*InvC-opsin WT in the presence of 1 μM retinal. Luminescence levels are normalized to the value at the starting point (time = 0 min). Error bars indicate the S.D. values (n = 3). Blue (+B) and orange (+O) bars indicate blue and orange illuminations, respectively. Before the measurements, intracellular cAMP levels were elevated by application of 1 μM forskolin. *C*, Representative trace showing orange and blue light-induced activation and deactivation of GIRK1/GIRK2 channels by *Acr*InvC-opsin WT in a *Xenopus* oocyte. Time lapse change of the current amplitude at -100 mV is plotted. Blue and orange light stimulations were applied at the times indicated by the blue and orange bars, respectively. Similar results were observed in independently injected oocytes and shown in Supplemental Fig. S5, A and B.

The NanoBiT assay clearly showed blue light-dependent Gi activation and orange light-dependent (partial) deactivation by *Acr*InvC-opsin. Thus, we examined the typical Gi/o-dependent cellular responses driven by the opsin activation. Upon Gi/o activation, dissociated Gi/oα inhibits adenylyl cyclase, which leads to a decrease in the intracellular cAMP levels, and dissociated Gβγ activates GIRK, which leads to an increase in the K^+^ currents^13,24^. We analyzed the changes in intracellular cAMP levels using the GloSensor cAMP assay^25^ in COS-1 cells and GIRK activation electrophysiologically using a two-electrode voltage clamp in *Xenopus* oocytes^10^. Our data from the GloSensor assay indicated that the first orange light illumination decreased cAMP levels, the second blue light illumination caused a further decrease, and the third orange light illumination partially terminated the light-induced cAMP reduction (Fig. 2B). The first orange light-induced cAMP reduction in the GloSensor assay (Fig. 2B) was consistent with the orange light-dependent Gi dissociation in the NanoBiT assay (Fig. 2A). Similarly, the first orange light stimulation caused significant GIRK activation in the *Xenopus* oocytes, and the second blue light stimulation increased GIRK current (Figs. 2C, and Supplemental Fig. S5, A and B). The first orange light-induced responses and further responses to the second blue light illumination were consistently observed in the NanoBiT, GloSensor, and GIRK measurements (Fig. 2). Also, in NanoBiT and Glosensor assays as well as GIRK current measurement, the third orange light illumination terminated the responses caused by the second blue light illumination, indicating that *Acr*InvC-opsin-induced responses can be reversibly modulated by blue and orange light (Fig. 2).

One possible explanation for the first orange light-induced cAMP and GIRK responses is that only a small amount of activated *Acr*InvC-opsin (see Supplemental Figs. S3 and S4) activates a substantial amount of Gi/o, leading to the downstream second messenger and ion channel responses (opsin to Gi/o to adenylyl cyclase or GIRK channel)^26^. In particular, the second messenger cAMP responses is amplified through the signal cascade. If so, a reduction in the number of functional *Acr*InvC-opsin in cells would severely affect the orange light-induced responses rather than blue light-induced responses. To test this possibility, we reduced the amount of functional *Acr*InvC-opsin in COS-1 cells and conducted a GloSensor assay using the same sequential light illuminations (orange-blue-orange-blue-orange). To achieve this, we reduced the amount of retinal added in the GloSensor assay (Fig. 3A) and the amount of opsin plasmid used for transfection (Fig. 3C). We compared the cAMP responses under the same sequential light illumination between the experimental conditions.

**Fig. 3.**
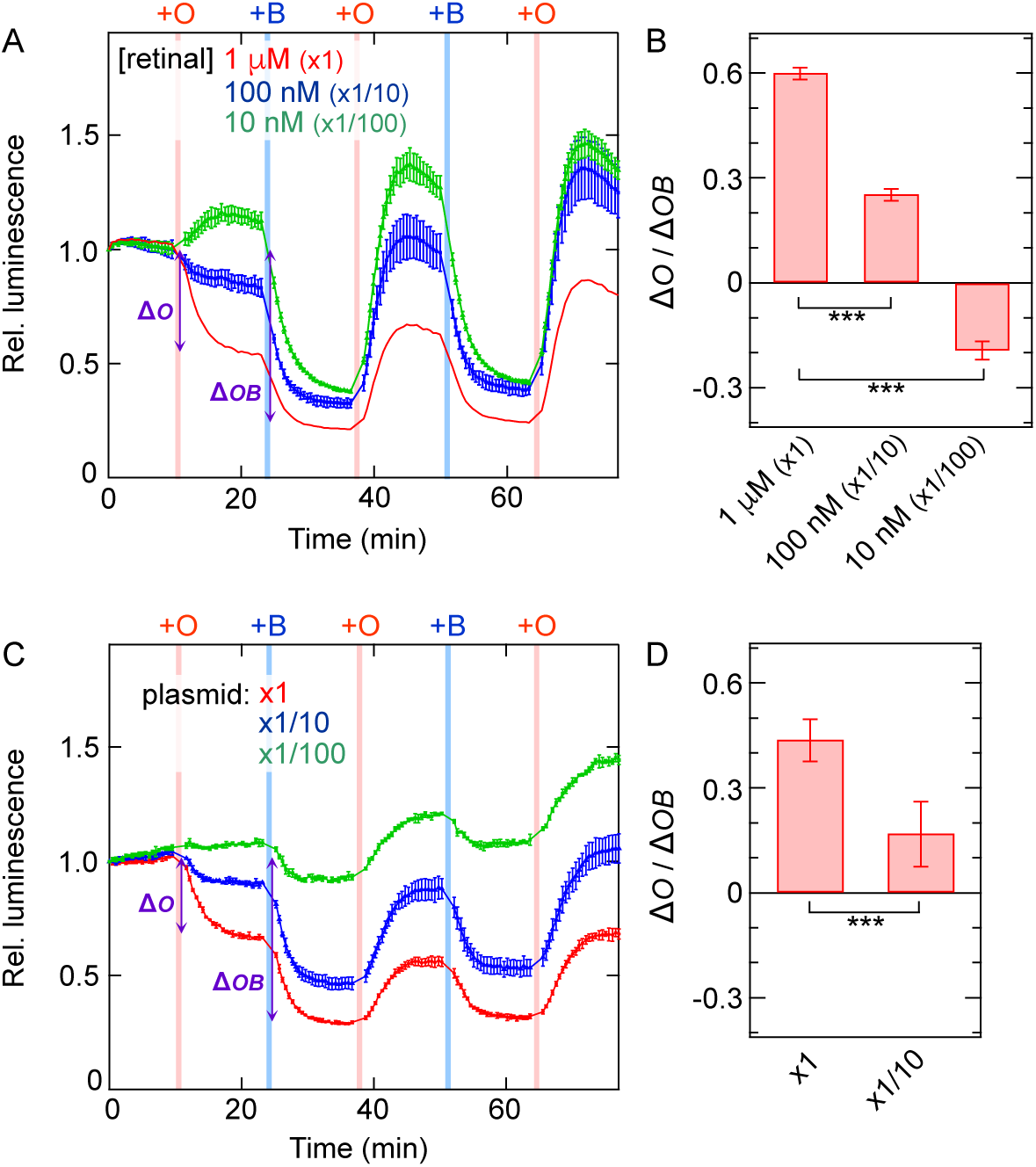
Light induced changes in GloSensor luminescence in COS-1 cells transfected with different amount of plasmid coding *Acr*InvC-opsin WT. *A*, Light induced changes in cAMP biosensor (GloSensor) luminescence in COS-1 cells expressing *Acr*InvC-opsin WT in the presence of different concentrations of exogenous retinal. Luminescence levels are normalized to the value at the starting point (time = 0 min). *Red, blue*, and *green* data points indicate relative luminescence levels in the presence of 1 μM (x1), 100 nM (x1/10), and 10 nM (x1/100) exogenous 11-*cis*-retinal, respectively. The data of *“*1 μM (x1)” (*red*) are adopted from Fig. 2B. Error bars indicate the S.D. values (n = 3). Blue (+B) and orange (+O) bars indicate blue and orange illuminations, respectively. Before the measurements, intracellular cAMP levels were elevated by application of 1 μM forskolin. *B*, Comparison of the ratio between luminescence changes upon the first orange light illumination (“β*O*” in panel *A*) and upon the first orange and the second blue light illuminations (“β*OB*” in panel *A*) in the presence of different amount of retinal. Error bars indicate the S.D. values (n = 3). The value of “1 μM (x1)” is significantly different from the values of “100 nM (x1/10)” and “10 nM (x1/100)” (*P* < 0.001***). The statistical differences were evaluated using Tukey’s test following one-way ANOVA. *C*, Luminescence levels are normalized to the value at the starting point (time = 0 min). *Red, blue*, and *green* data points indicate relative luminescence levels of COS-1 cells transfected with x1 (50 ng/well), x1/10 (5 ng/well), x1/100 (0.5 ng/well) amount of *Acr*InvC-opsin WT plasmids. Before the measurements, 1 μM 11-*cis*-retinal was added and intracellular cAMP levels were elevated by application of 1 μM forskolin. Error bars indicate the S.D. values (n = 3). Blue (+B) and orange (+O) bars indicate blue and orange illuminations, respectively. Since under the “x1/100” conditions, even blue light illumination (+B) induced small cAMP responses, the “1/100” data were excluded from the statistical analyses in panel *D*. *D*, Comparison of the ratio between luminescence changes upon the first orange light illumination (“β*O*” in panel *C*) and upon the first orange and the second blue light illuminations (“β*OB*” in panel *C*) in COS-1 cells transfected with different amount of *Acr*InvC-opsin coding plasmid. Error bars indicate the S.D. values (n = 6). The value of “x1” is significantly different from the value of “x1/10” (*P* < 0.001*** by Student’s unpaired t-test).

### Light-induced cellular responses in the presence of reduced amount of functional AcrInvC-opsin molecules

We conducted the GloSensor assays on *Acr*InvC-opsin with 1 μM (x1), 100 nM (x1/10), or 10 nM (x1/100) retinal in order to compare light-dependent cAMP responses in the presence of different amount of functional *Acr*InvC-opsin molecules. Under the “100 nM (x1/10)” conditions, the first orange light illumination caused a smaller cAMP reduction than under the “1 μM (x1)” conditions, but the second blue light decreased the cAMP levels to a similar extent under both the conditions (Fig. 3, A and B). The ratio of luminescence changed upon illumination with the first orange light (“*ΔO*” in Fig. 3A) relative to the sum of changes upon illumination with the first orange and the second blue light (“*ΔOB*” in Fig. 3A) was significantly different between the two conditions (Fig. 3B). Under the “10 nM (x1/100)” conditions, the first orange light caused a slight increase in cAMP levels (Fig. 3, A and B). This was probably because some of the added 11-*cis*-retinal was thermally isomerized to the all-*trans* isomer in solution, and under retinal-depleted (opsin-excess) conditions, some *Acr*InvC-opsin proteins bound to the isomerized all-*trans*-retinal to form the activated state. Orange light deactivates the all-*trans*-retinal bound (activated) opsin molecules, leading to the slight increase in cAMP levels. The second blue light illumination caused a robust cAMP reduction similar to the other conditions (Fig. 3B).

Many functional (light-sensitive) opsin molecules are formed in the presence of sufficient retinal molecules. Upon orange light illumination, a very small proportion was activated, but the total number of activated molecules would be large enough to induce a substantial cAMP response (Fig. 2B). In contrast, after addition of limited amount of retinal, the total number of functional opsin molecules reduced. Upon orange light illumination, a small proportion of the reduced number of opsin molecules was activated, but this was not sufficient to induce large responses (Fig. 3, A and B). Even under retinal depletion conditions, blue light illumination can convert a large proportion of opsin molecules into an activated state, leading to substantial cellular responses that are amplified in the signaling cascade (opsin to Gi/o to adenylyl cyclase).

Similarly, the GloSensor assay on COS-1 cells transfected with “x1/10” amount of plasmid for *Acr*InvC-opsin showed slight response to the first orange light-induction but the second blue light induced substantial cAMP responses (Fig. 3, C and D). In this case, the expression level of opsin protein was reduced, and the number of activated opsin molecules upon orange light illumination was too small to induce substantial cAMP responses. Under the “x1/100” plasmid conditions, the orange light-induced cAMP responses were negligible, and even the blue light-induced responses largely decreased (Fig. 3C), suggesting that total activated opsin molecules were insufficient to evoke cellular responses even after blue light illumination. Thus, our data using reduced amount of retinal or *Acr*InvC-opsin-coding plasmid, “appropriate” amount of functional opsin molecules in cells would enable to minimize unintentional orange light-induced cellular responses while maintaining substantial blue light-indued responses (see “x1/10” condition data in Fig. 3, A and C).

### Modulation of molecular properties of AcrInvC-opsin by an amino acid substitution

Our spectral and signaling analyses of *Acr*InvC-opsin WT revealed that this opsin is a visible light-sensitive bistable pigment in which the resting and activated states can be inter-converted by blue and orange light illumination (Figs. 1 and 2). Activated *Acr*InvC-opsin can drive typical Gi/o-dependent signaling pathways in a sustained manner. Similar to other opsins, *Acr*InvC-opsin possessed broad absorption spectra (Fig. 1B, and Supplemental Fig. S4), and orange light illumination of the opsin leads to the formation of a tiny amount of the activated state (see Supplemental Fig. S3), resulting in substantial downstream cellular responses under certain experimental conditions (Fig. 2). The unintentional orange light-induced activation could be due to the spectral overlap between the resting and activated states (Supplemental Fig. S4). We attempted to decrease the spectral overlap by amino acid substitution to minimize orange light-induced activation. Compared to the abundant information of spectral tuning mechanisms in vertebrate visual pigments, the mechanisms of InvC-opsins are far less understood. It is also unknown how the spectrum of activated opsin is tuned (but see refs 27, 28). In our previous study, we identified an amino acid residue at position 94, which is close to the Schiff base linkage with the retinal, is responsible for spectral tuning in *Platynereis* c-opsin1^10^. *Acr*InvC-opsin contains a serine residue at this position (Fig. 4A and Supplemental Fig. S2A), and we noticed the Ser-94 residue to potentially play an important role in regulating the spectral properites^29^. Based on these insights, we introduced the S94A substitution into *Acr*InvC-opsin, and analyzed the spectral properties of the purified mutant protein.

**Fig. 4.**
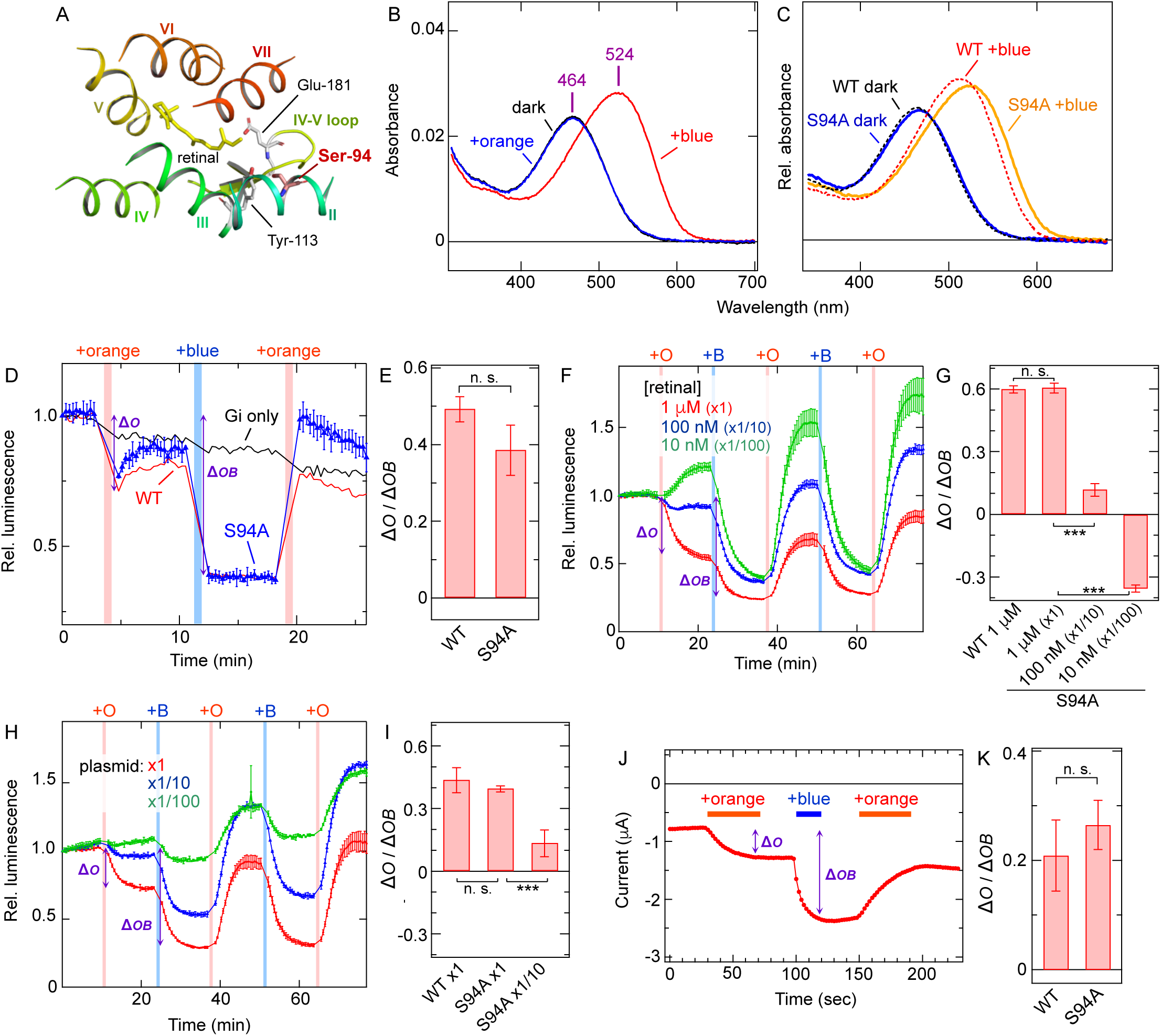
Spectral and signaling properties of *Acr*InvC-opsin S94A mutant. *A*, Arrangement of Ser-94, Tyr-113, and Glu-181 in the AlphaFold2-predicted structure^34^ of *Acr*InvC-opsin WT. Glu-181 is a putative counterion for the protonated Schiff base linkage that is important for visible light absorption. Amino acid at position 113 (occupied with Glu in vertebrate visual pigment) is the counterion for he protonated Schiff base linkage in vertebrate visual pigments. The 11-*cis*-retinal molecule is adopted from the crystal structure of bovine rhodopsin (PDB ID: 1U19)^35^. The structural models are prepared using PyMOL (https://pymol.org/). *B*, Absorption spectra of purified S94A mutant of *Acr*InvC-opsin. Spectra in the dark (*black*), after blue illumination (*red*), and after subsequent orange light (>580 nm) illumination (*blue*) are shown. In the dark, α_max_ value is 464 nm, and after blue light illumination, the value is shifted to 524 nm. *C*, Comparison of absorption spectra between *Acr*InvC-opsin WT and S94A mutant. Normalized absorption spectra of *Acr*InvC-opsin WT in the dark (*black*, dotted) and after blue light absorption (*red*, dotted), and S94A mutant in the dark (*blue*) and after blue light absorption (*yellow*) are shown. Spectra of WT and S94 are normalized based on absorbance values in the dark. *D*, NanoLuc luminescence changes of NanoBiT-tagged Gi proteins upon blue and orange light illumination on *Acr*InvC-opsin S94A mutant (*blue*) in COS-1 cells. The data of *Acr*InvC-opsin WT (*red*) and Gi only (*black*) are adopted from Fig. 2A. Error bars indicate the S.D. values (n = 3). NanoLuc luminescence levels are normalized to the value at the starting point (time = 0 min). *E*, Comparison of the ratio between luminescence changes upon the first orange light illumination (“β*O*” in panel *D*) and upon the first orange and the second blue light illuminations (“β*OB*” in panel *D*) of *Acr*InvC-opsin WT and S94A. Error bars indicate the S.D. values (n = 3). The “β*O* / β*OB*” values were not significantly different between WT and the S94A mutant (*P* = 0.064 by Student’s unpaired t-test). *F*, Light induced changes in cAMP biosensor (GloSensor) luminescence in COS-1 cells expressing *Acr*InvC-opsin S94A in the presence of different concentrations of exogenous retinal. Luminescence levels are normalized to the value at the starting point (time = 0 min). *Red, blue*, and *green* data points indicate relative luminescence levels in the presence of 1 μM (x1), 100 nM (x1/10), and 10 nM (x1/100) exogenous 11-*cis*-retinal, respectively. Before the measurements, intracellular cAMP levels were elevated by application of 1 μM forskolin. Error bars indicate the S.D. values (n = 3). *G*, Comparison of the ratio between luminescence changes upon the first orange light illumination (“β*O*” in panel *F*) and upon the first orange and the second blue light illuminations (“β*OB*” in panel *F*) in the presence of different amount of retinal. Error bars indicate the S.D. values (n = 3). The “WT 1 μM” value is adopted from Fig. 3B, and this value is not significantly different from the “S94A 1 μM (x1)” value (*P* = 0.988). The value of “S94A 1 μM (x1)” is significantly different from the values of “S94A 100 nM (x1/10)” and “S94A 10 nM (x1/100)” (*P* < 0.001***). The statistical differences were evaluated using Tukey’s test following one-way ANOVA. *H*, Light induced changes in GloSensor luminescence in COS-1 cells transfected with different amount of plasmid coding *Acr*InvC-opsin S94A mutant. Luminescence levels are normalized to the value at the starting point (time = 0 min). *Red, blue*, and *green* data points indicate relative luminescence levels of COS-1 cells transfected with x1 (50 ng/well), x1/10 (5 ng/well), x1/100 (0.5 ng/well) amount of the S94A mutant plasmids, respectively. Before the measurements, 1 μM 11-*cis*-retinal was added and intracellular cAMP levels were elevated by application of 1 μM forskolin. Error bars indicate the S.D. values (n = 3). Since under the “x1/100” conditions, even blue light illumination (+B) induced small cAMP responses, the “1/100” data were excluded from the statistical analyses in panel *I*. *I*, Comparison of the ratio between luminescence changes upon the first orange light illumination (“β*O*” in panel *H*) and upon the first orange and the second blue light illuminations (“β*OB*” in panel *H*) in COS-1 cells transfected with different amount of *Acr*InvC-opsin coding plasmid. Error bars indicate the S.D. values (n = 6). The “WT x1” value is adopted from Fig. 3D, and this value is not significantly different from the “S94A x1” value (*P* = 0.359). The value of “S94A x1” is significantly different from the value of “S94A x1/10” (*P* < 0.001***). The statistical differences were evaluated using Tukey’s test following one-way ANOVA. *J*, Representative trace showing orange and blue light-induced activation and deactivation of GIRK1/GIRK2 channels by *Acr*InvC-opsin S94A mutant in a *Xenopus* oocyte. Time lapse change of the current amplitude at -100 mV is plotted. Blue and orange light stimulations were applied at the times indicated by the blue and orange bars, respectively. Similar results were observed in independently injected oocytes and shown in Supplemental Fig. S5, C and D. *K*, Comparison of the ratio between luminescence changes upon the first orange light illumination (“β*O*” in panel *J*) and upon the first orange and the second blue light illuminations (“β*OB*” in panel *K*) of *Acr*InvC-opsin WT and S94A. Error bars indicate the S.D. values (n = 8 for WT and n = 9 for the S94A mutant). The “β*O* / β*OB*” values were not significantly different between WT and the S94A mutant (*P* = 0.054 by Student’s unpaired t-test). In panels *D*, *F, H,* and *J*, blue (+blue or +B) and orange (+orange or +O) bars indicate blue and orange illuminations, respectively.

The purified *Acr*InvC-opsin S94A mutant showed αmax at 464 nm (Fig. 4B), which was almost identical to αmax of WT (Figs. 1B and 4C). This result indicates that the Ser-94 does not play an important role in the spectral tuning of the resting state. Upon blue light illumination, the S94A mutant showed a red-shifted spectrum with αmax at 524 nm (Fig. 4, B and C), indicating that this residue plays a role in the spectral tuning of the activated state. This spectral shift is consistent with a previous study showing that introduction of the second transmembrane region (including position 94) of *Drosophila* Rh2 into *Drosophila* Rh1 caused a spectral shift in the activated state of Rh1^28^. The spectral shift specific in the active state indicated that the spectral overlap between the resting and active states decreased by the substitution (Fig. 4C). The red-shift in the activated state would make the state more efficiently absorb orange light, potentially leading to the suppression of unintentional orange light-induced activation. To examine this effect, we conducted NanoBiT Gi dissociation assay, GloSensor assay, and GIRK current measurements on the S94A mutant.

In NanoBiT assay on the S94A mutant, the first orange illumination caused a slight smaller Gi dissociation, but the ratio of luminescence changed upon the first orange light (“*ΔO*” in Fig. 4D) relative to the sum of changes upon the first orange and the second blue lights (“*ΔOB*” in Fig. 4D) was not significantly different between the S94A mutant and WT (Fig. 4E). In addition, the GloSensor assay and GIRK current measurement of the S94A mutant showed that the first orange light illumination of the S94A mutant caused substantial cAMP responses and GIRK activation, as was observed with respect to WT (Fig. 4, F and J, and Supplemental Fig. S5, C and D). In both analyses, the ratios of changes upon the first orange light (“*ΔO*” in Fig. 4, F and J) relative to the sums of changes upon the first orange and the second blue lights (“*ΔOB*” in Fig. 4, F and J) were not significantly different between the S94A mutant and WT (Fig. 4, G and K, and Supplemental Fig. S5, C and D). Thus, the activated state-specific ∼10 nm spectral shift caused by the S94A substitution was not sufficient to suppress the unintentional orange light-induced responses. In other words, a 60 nm spectral separation between the resting and activated states would be insufficient to complete deactivation by selective illumination of the activated state.

We also investigated the GloSensor assay profiles of the S94A mutant in the presence of a smaller amount of functional opsin molecules by changing the amount of added retinal molecules (Fig. 4F) or the opsin mutant-coding plasmid (Fig. 4H). As observed with respect to WT (Fig. 3, B and D), the ratio of “*βO*/*βOB*” was significantly decreased under conditions of “100 nM (x1/10)” retinal or “1/10” plasmid (Fig. 4, G and I). These results indicate that “appropriate” amounts of the S94A *Acr*InvC-opsin molecules can minimize unintentional orange light-induced cellular responses while maintaining substantial blue light-indued responses. In addition, under “10 nM (x1/100)” retinal and “x1/100” plasmid conditions, the S94A mutant showed similar profiles with respect to WT as well (Figs, 3A, 3C, 4F, and 4H). Thus, the mutation-induced ∼10 nm spectra separation was not sufficient to alter the light-induced second messenger responses.

## Discussion

In the present study, we characterized *Acr*InvC-opsin as a blue-sensitive, Gi/o-coupled, and bistable pigment. The activation and deactivation of the opsin can be bidirectionally regulated by illumination with blue or orange light. Mutational analyses of *Acr*InvC-opsin clearly indicated that the spectral properties of the activated state could be selectively tuned by the S94A substitution. These molecular characteristics of *Acr*InvC-opsin would help understand the photoreceptive functions in the fan worm *Acromegalomma interruptum* and closely related animals. In addition, the molecular properties would be useful for engineering a new optical control tool to regulate Gi/o-biased signalings using different colors of visible light.

### Relationship of molecular characteristics of AcrInvC-opsin and photoreceptive functions in Acromegalomma interruptum

A previous study on the fan worm *Acromegalomma interruptum* identified *Acr*InvC-opsin as a putative photoreceptive protein in the radiolar eyes^16^, but functional properties of the opsin have not been tested (see “Introduction”). In a subsequent study, microspectrophotometry was conducted on radiolar eyes in a closely related annelid *Spirobranchus corniculatus*, and the eyes showed αmax at 464 nm^29^. The αmax at 464 nm of the resting state of *Acr*InvC-opsin (Fig. 1B) was consistent with the previous studies. These insights and our present study strongly suggest that the InvC-opsin functions as a blue light receptor in the radiolar eyes, and blue-sensitivity of radiolar eyes might be conserved among closely related annelids.

Our study also showed that *Acr*InvC-opsin is selectively coupled with Gi/o-type G proteins (Fig. 1, C and D) and that the opsin is a typical Gi/o-coupled bistable pigment wherein the resting and activated states are inter-convertible upon blue and orange light absorption (Fig. 1B). The activated state sustainedly activates Gi/o-type G proteins to evoke Gi/o-dependent cellular responses (Fig. 2). These results suggest that in the radiolar eyes of *Acromegalomma interruptum*, Gi/o-dependent signaling cascades are driven upon blue light reception. The previous study reported that Goα was highly expressed in the radiolar eyes based on transcriptome data^16^. Other InvC-opsins, such as *Platynereis* c-opsin1/2 and mosquito Opn3, have been reported to drive Gi/o signaling pathways in a light-dependent manner and play important physiological roles in the animals^4,7,9,11,12,30,31^. Photoreceptive functions involving Gi/o-coupled and bistable InvC-opsins will be extensively investigated to understand visual and non-visual photoreceptions in various invertebrates.

### Potentials of AcrInvC-opsin as a Gi/o-biased optical control tool

In recent optogenetic studies, Gi/o-coupled bistable opsins have been used as inhibitory optical control tools to suppress cellular activity through intracellular cAMP reduction and/or GIRK activation^8,14,15,32,33^. InvC-opsins such as mosquito Opn3 and *Platynereis* c-opsin1 have been utilized as efficient optical control tools to suppress neural activities by light^13–15^. As mentioned in “Introduction”, mosquito Opn3 cannot be deactivated by light because of its highly spectral overlap, and *Platynereis* c-opsin1 is UV-sensitive. *Acr*InvC-opsin could be useful as a bistable optical control tool that is selectively coupled with Gi/o (Figs. 1C and 2) and can be turned ON and OFF by visible light (Figs. 2). On the other hand, our data showed that in the presence of a large number of photoreceptive opsin molecules, deactivating orange light unintentionally leads to substantial cellular responses (Figs. 2 and 3), owing to a slight overlap between the absorption spectrum of the resting state and the radiation spectrum of orange light (Supplemental Figs. S3 and S4). A similar unintentional cellular response to deactivating light was reported in optogenetics using lamprey parapinopsin, which is UV-sensitive and is deactivated by visible light. Blue light illumination on parapinopsin caused substantial (but less than UV-dependent) neural responses^32^. This unintentional activation is primarily due to spectral broadness and overlap in opsins.

The S94A substitution-induced selective spectral shift in the activated state and a decrease in the spectral overlap between the resting and activated states of *Acr*InvC-opsin (Fig. 4C) were not sufficient to solve this problem as an optical control tool (Fig. 4, D – K). If the functional opsin number is too high in the target cells, the deactivating orange light would induce unintentional activating responses. On the other hand, when the number of functional opsins was too low, the activating blue light could not induce sufficient light-dependent responses (see “1/100” plasmid conditions in Figs. 3C and 4H). “Appropriate” expression levels of *Acr*InvC-opsin in the target cells will enable clear ON/OFF responses to blue and orange light illuminations. However, in typical optogenetic analyses, fine control of functional opsin number in tissues is not feasible. The spectral shift cause by the S94A substitution proved that the absorption spectra of the resting or activated states can be independently changed (Fig. 4, B and C). Additional substitution(s) could further decrease the spectral overlap to completely diminish the unintentional opsin activation by deactivating light. Thus, *Acr*InvC-opsin has the potential to act as a Gi/o-biased optical control tool that can be activated and deactivated by visible light.

## Conclusions

Characterization of the spectral and signaling properties of *Acr*InvC-opsin provided important insights into the photoreceptive functions in the fan worm and related annelids. Furthermore, the bistable nature of *Acr*InvC-opsin, wherein the resting and activated states can be inter-convertible by visible light illumination, is important for constructing useful Gi/o-biased optical control tools.

## Author contributions

S. F., T. K., and H. T. designed the study, conducted experiments, and analyzed obtained data. T. S. and Y. K. designed and supervised electrophysiological experiments. All authors discussed obtained data and wrote the paper.

## Competing interests

The authors declare no competing interests.

## Acknowledgment

We thank Dr. David Farrens (Oregon Health & Science University) for providing us with COS-1 cell line and the pMT expression vector, and Dr. Robert Molday (University of British Columbia, Canada) for providing hybridoma cells producing 1D4 antibody. We also thank Ms. Tomomi Yamamoto, Ms. Chizue Naito (National Institute for Physiological Science), Ms. Hiroe Motomura, Ms. Kayo Inaba (Institute for Molecular Science), and the Functional Genomics Facility, NIBB Core Research Facilities (Okazaki, Japan) for technical support.

## Funding

H. T. is supported by JST, PRESTO (JPMJPR1787), the Japan Society for the Promotion of Science KAKENHI Grant 21H02445, and the Center for the Promotion of Integrated Sciences of SOKENDAI. T. S. is supported by the Japan Society for the Promotion of Science KAKENHI Grant 23K06352. Y. K. is supported by the Japan Society for the Promotion of Science KAKENHI Grant JP23H02667. This study was supported by the Cooperative Study Program (19-254, 20-267, 21-264, 22NIPS239, 23NIPS104, 24NIPS131) of National Institute for Physiological Sciences.

## Abbreviations

c-opsin: ciliary opsin
Gi/o: Gi/o-type trimeric G proteins
GIRK: G-protein-coupled inwardly rectifying potassium channel
GPCR: G protein-coupled receptor
*Acr*InvC-opsin: *Acromegalomma* invertebrate ciliary opsin
αmax: absorption maximum
UV: ultra-violet
WT: wild-type

## Supporting Information

**Supplemental Fig. S1.**
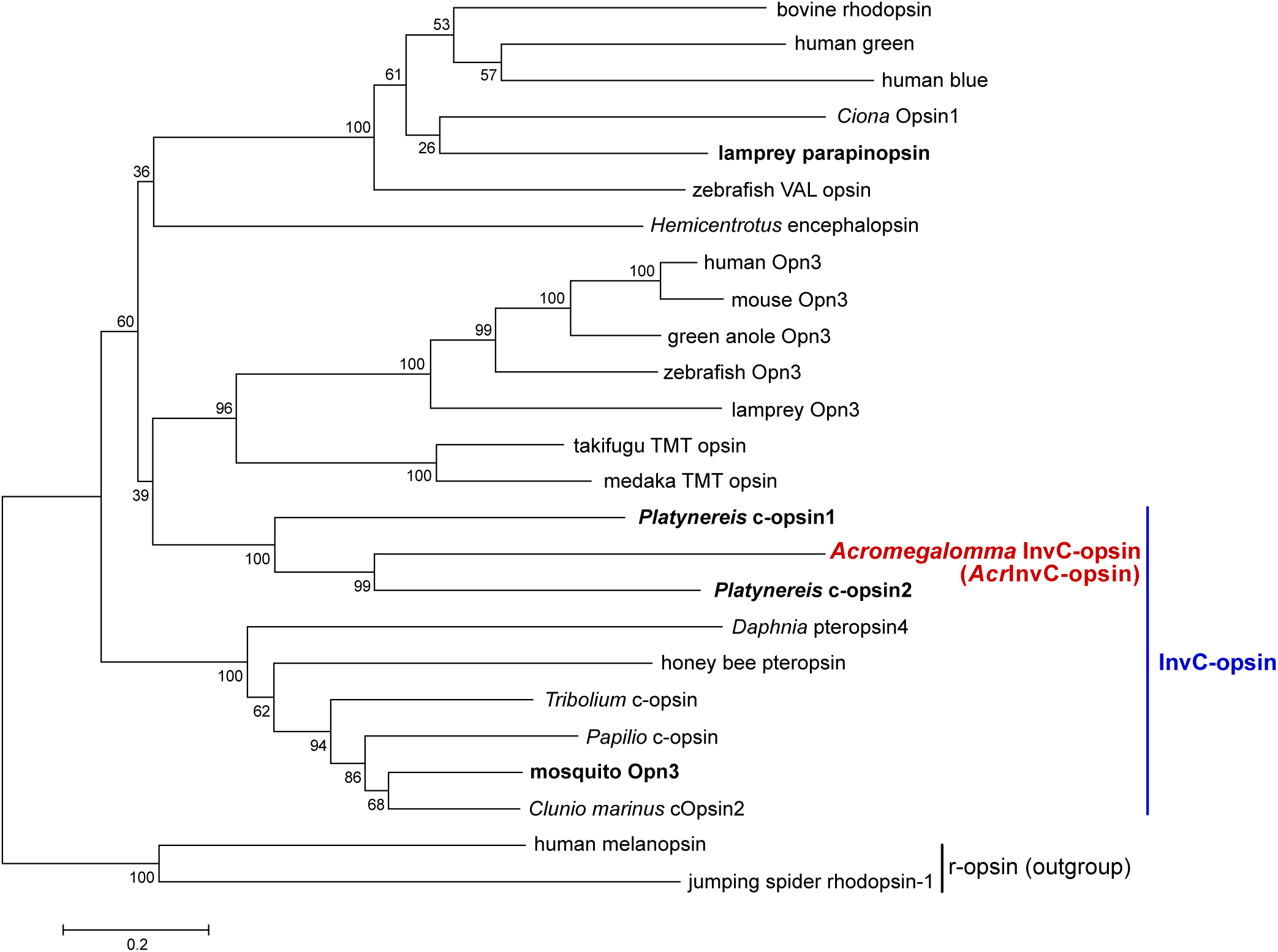
Molecular phylogenetic tree of ciliary opsins. Molecular phylogenetic relationship of *Acr*InvC-opsin and other ciliary opsins is shown. The phylogenetic tree was constructed by the neighbor-joining method using MEGA10 software^36^. InvC-opsin group and r-opsin (rhabdomeric opsin) outgroup are indicated. Ciliary opsins mentioned in the main text are highlighted in boldface. The bootstrap probabilities are indicated at each branch node, and the scale bar (0.2 substitutions per site) is also shown.

**Supplemental Fig. S2.**
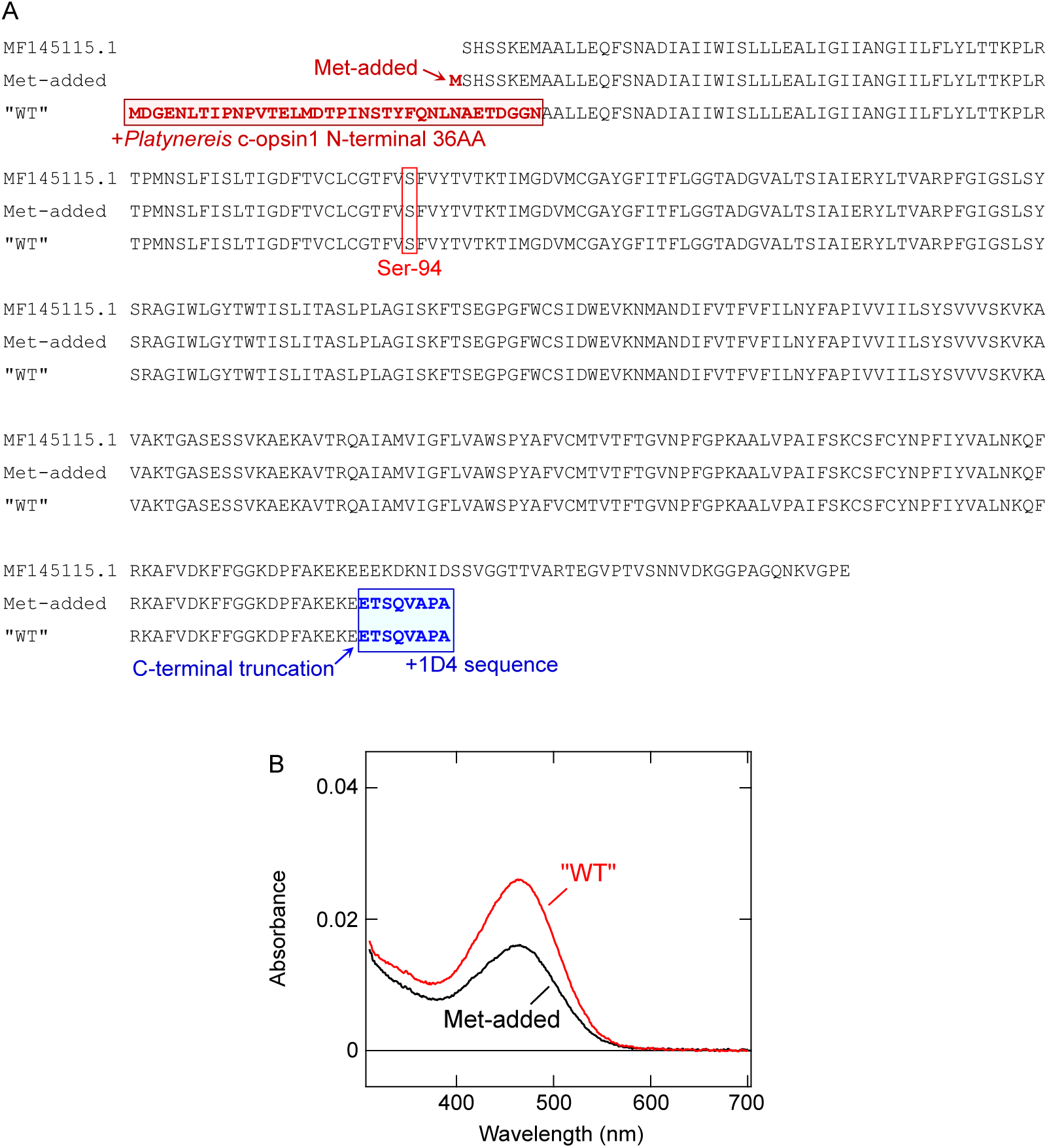
The construct of *Acr*InvC-opsin used in this study. *A*, Amino acid sequence comparison among the reported partial sequence of *Acr*InvC-opsin (MF145115.1)^16^, addition of the initiation methionine with truncation of C-terminus as well as addition of the 1D4 sequence (“Met-added”), and addition of *Platynereis* c-opsin1 N-terminal 36 amino acids with truncation of C-terminus as well as addition of the 1D4 sequence (“WT”). Modified amino acids are colored and highlighted in boldface. The position of Ser-94 is also indicated. *B*, Comparison of expression levels between “Met-added” (*black*) and “WT” (*red*) constructs. Both constructs were expressed in the same number (10 plates) of dishes and purified in parallel. The result indicates that addition of *Platynereis* c-opsin1 N-terminal 36 amino acids on *Acr*InvC-opsin increased expression levels in comparison with the simple addition of the initiation methionine.

**Supplemental Fig. S3.**
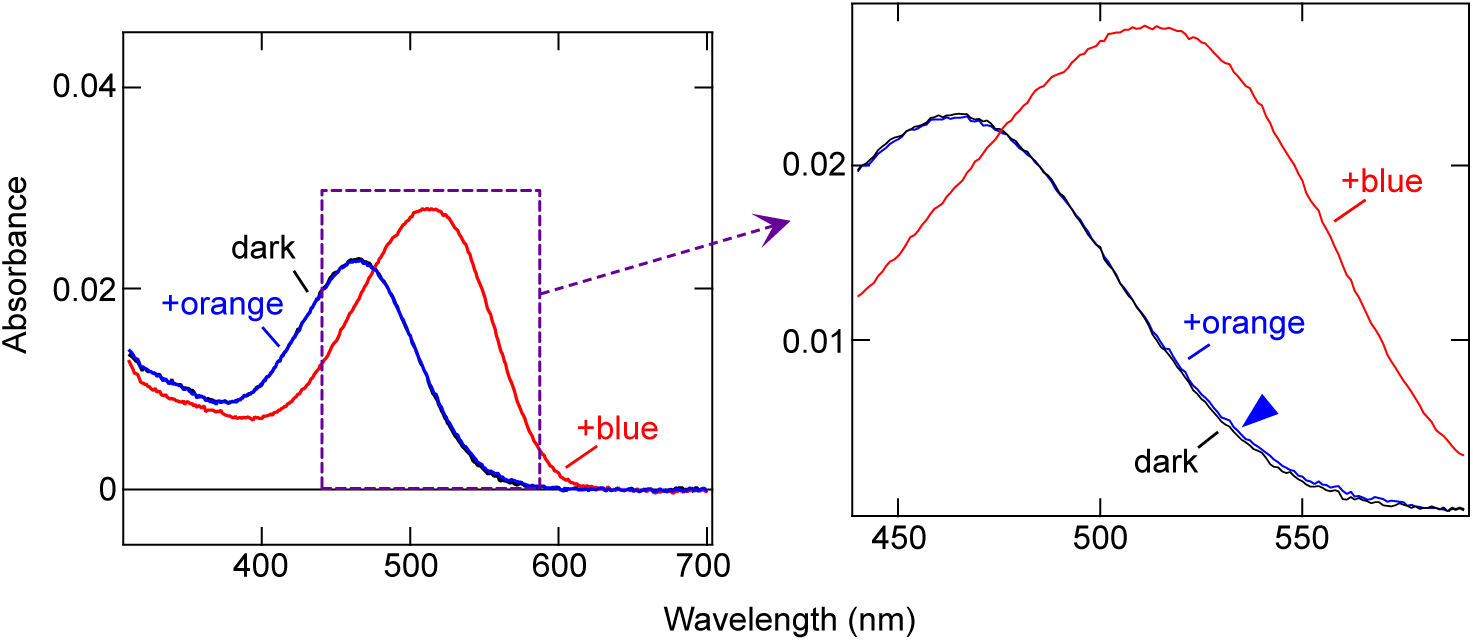
Very tiny amount of the red-shifted species of *Acr*InvC-opsin after orange light illumination. The absorption spectra of *Acr*InvC-opsin WT are adopted from Fig. 1B (*left*), and the highlighted region (purple dotted square) is expanded (*right*). After orange light illumination, very tiny amount of the red-shifted species (*blue arrowhead*) was remained. See main text for details.

**Supplemental Fig. S4.**
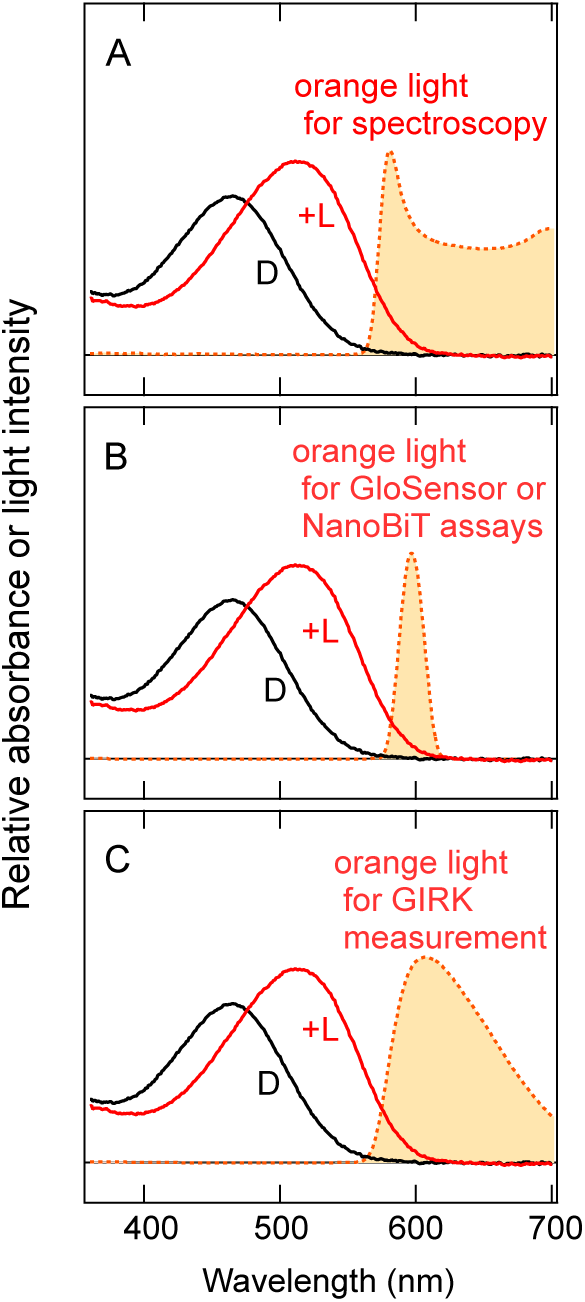
Spectral overlap of absorption spectra of *Acr*InvC-opsin with radiation spectra of deactivating “orange” lights used for various assays. *A, B, C,* Normalized absorption spectra of *Acr*InvC-opsin WT in the dark (black line, “D”) and after blue light absorption (red line, “+L”) and radiation spectra of orange lights (light orange shaded area with dotted orange line) used for spectroscopy (*A*), GloSensor and NanoBiT assays (*B*), and GIRK measurement (*C*) are shown. The orange lights are primarily absorbed by the activated state (“+L”) due to the large spectral overlap, but much less efficiently absorbed by the resting state (“D”) due to the tiny overlap.

**Supplemental Fig. S5.**
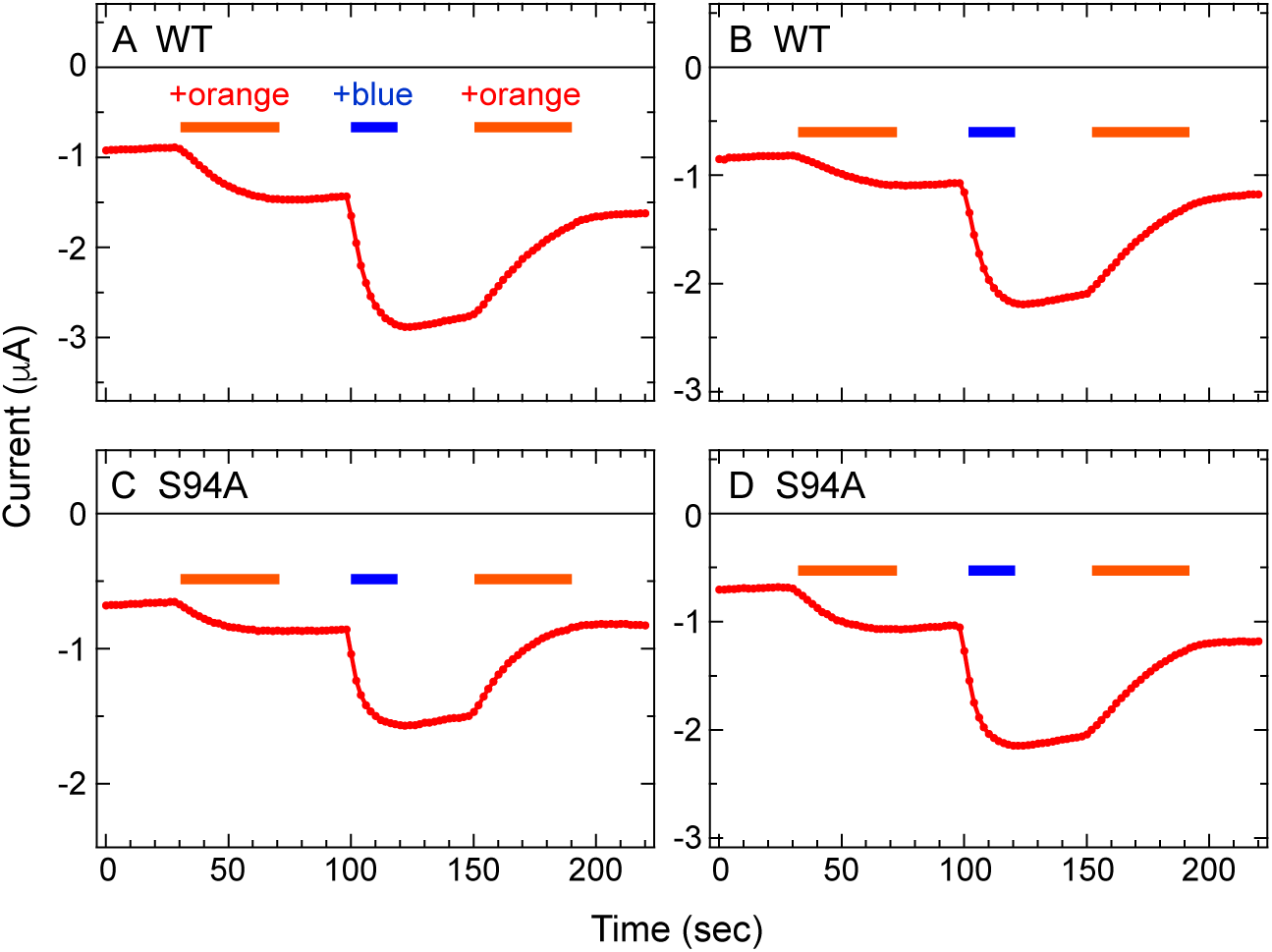
Orange and blue light-induced activation and deactivation of GIRK1/GIRK2 channels by *Acr*InvC-opsin WT and S94A mutant in *Xenopus* oocytes. GIRK current trances in *Xenopus* oocytes expressing *Acr*InvC-opsin WT (*A, B*) or S94A mutant (*C, D*) are shown. Time lapse changes of the current amplitude at -100 mV are plotted. Blue and orange light stimulations were applied at the times indicated by the blue and orange bars, respectively.

## Notes

### Competing Interest Statement

The authors have declared no competing interest.

